# Identifiability and inference of phylogenetic birth-death models

**DOI:** 10.1101/2022.08.26.505438

**Authors:** Brandon Legried, Jonathan Terhorst

## Abstract

Recent theoretical work on phylogenetic birth-death models offers differing viewpoints on whether they can be estimated using lineage-through-time data. Louca and Pennell (2020) showed that the class of models with continuously differentiable rate functions is nonidentifiable: any such model is consistent with an infinite collection of alternative models, which are statistically indistinguishable regardless of how much data are collected. Legried and Terhorst (2022a) qualified this grave result by showing that identifiability is restored if only piecewise constant rate functions are considered.

Here, we contribute new theoretical results to this discussion, in both the positive and negative directions. Our main result is to prove that models based on piecewise polynomial rate functions of any order and with any (finite) number of pieces are statistically identifiable. In particular, this implies that spline-based models with an arbitrary number of knots are identifiable. The proof is simple and self-contained, relying mainly on basic algebra. We complement this positive result with a negative one, which shows that even when identifiability holds, rate function estimation is still a difficult problem. To illustrate this, we prove some rates-of-convergence results for hypothesis testing using birth-death models. These results are information-theoretic lower bounds which apply to all potential estimators.

## 1 Introduction

The linear birth-death (BD) process (Feller, 1939; Kendall,1948) has been used to study population growth in a variety of settings. Recently, in phylogenetics, it has also served as a model of tree formation, by viewing the surviving lineages of a tree as members of population, which randomly give birth to other lineages, or go extinct. Often, the rates at which these “births” and “deaths” occur in a phylogeny touch on important evolutionary questions. For example Nee et al. (1994); Quental and Marshall (2010); Morlon et al. (2011) used phylogenetic BD models to study extinction and speciation dynamics; Gernhard (2008); Heath et al. (2014) used them to calibrate divergence times; and Stadler (2009, 2010); Stadler et al. (2013) investigated the dynamics of pathogens in an infection tree. Estimating the rate at which lineages are born and die in an observed phylogeny is not trivial: in the usual case where only extant members of the population can be sampled, deaths are not recorded at all, and the apparent rate of births will also be biased downwards, because some lineages went extinct before the present, or were simply not sampled. The birth-death model of tree formation provides a principled way to correct these biases.

Despite its widespread use, serious questions have recently been raised about whether it is even possible to estimate this model from phylogenetic data.Stadler(2009) showed that even when the birth and death rates are assumed constant over time, the birth-death model with present-day sampling fails to be identifiable if the sampling probability is not known in advance. That is, multiple distinct models produce exactly the same distribution over the observable data. Other studies have incorporated time-varying per capita birth and death rates in attempting to be more biologically realistic (Stadler et al., 2013). However, Louca and Pennell (2020) have recently shown that birth-death models with smoothly varying rate functions are unidentifiable from extant timetrees, meaning they cannot be reliably estimated using any amount of data.

In Legried and Terhorst (2022a), this grave finding was partly qualified by showing that a smaller class of candidate models, consisting of birth and death rates which are piecewise constant, is identifiable given a sufficiently large timetree. This sample size is small enough for many applications, such as the phylodynamic analysis of pathogens. A potential objection is that the class of piecewise-constant models considered by Legried and Terhorst (2022a) might not be sufficiently large if one believes that the birth and death rates could be continuous or satisfy other smoothness criteria. A partial response to this issue is given in their Theorem 5. Broadly speaking, that result states that any unidentifiable, but reasonably smooth, model can be approximated by an identifiable one. However, there is a sampling size requirement to have provable identifiability that diverges as the approximation error goes to zero. In their Conjecture 6, they suggest that their results extend to models with piecewise-polynomial (or spline) birth and death rates, with a similar sampling requirement to the piecewise-constant case. Resolution of this conjecture would give a sampling requirement for polynomial birth and death rates to be identifiable, but mathematical difficulty left the question open.

In this paper, we prove the following stronger version of their Conjecture 6: piecewise-polynomial models defined on an arbitrary (but finite) number of pieces are identifiable from extant timetree data. Further, the sample size requirement is removed. Our results demonstrate that it is *not* the presence of jump discontinuities that leads to identifiability. Indeed, our main result (Theorem 1) establishes the existence of identifiable model classes containing smooth (*C*^∞^) birth and death rate functions. The proof relies on the fact that polynomials, uniquely and by definition, have finite power series. In contrast, the proof technique of Legried and Terhorst (2022a) depends on solving a certain differential equation satisfied by the rate functions, which is difficult to carry out when those functions are not constant.

Identifiability is a minimal regularity criterion which must hold in order for a statistical model to be useful at all. Even when it does, the condition does not say anything about finite sample estimability, and indeed it often happens that identifiable models are quite difficult to estimate accurately. In the second section of the paper, we use ideas from a related literature on the estimation of coalescent-based models (Kim et al., 2015; Legried and Terhorst, 2022b) to study hypothesis testing problems for phylogenetic birth-death models. We derive information theoretic lower bounds which extend to all potential estimators of these models. Moreover, as these results assume perfect knowledge of the underlying time-tree, the problem could in fact be much harder given limited or error-prone data.

## 2 Background

Given *N* ≥ 2 samples, an *extant time tree* is a phylogenetic tree representing their ancestral relationships. The branching times are denoted *τ*_1_ > … >*τ*_*N*–1_ > 0, where time run backwards from the present. All *N* leaves are taken to be sampled at the present t = 0, so that the tree is ultrametric. There is also a source node called the *origin*, whose age is denoted by *τ_o_*, which is a user-specified parameter. Extant time trees are generated according to a phylogenetic birth-death (BD) process (Nee et al., 1994). The dynamics of this process are governed by three parameters. The per-capita birth and death rate functions 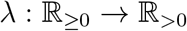 and 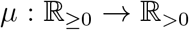 give the instantaneous rates at which surviving lineages give birth (bifurcate) or die. The third parameter *ρ* ∈ (0, 1] is the sampling probability; each lineage that survives to time *t* = 0 is sampled (exists in the time tree) with probability *ρ*, independently of all other surviving lineages. The triple (λ, *μ, ρ*) determines a phylogenetic BD process.

The birth and death rate functions are assumed to belong to a certain function space. In this paper, we study the case where they are given by piecewise-polynomial functions over the interval [0, *τ_o_*]. Let 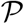 denote the set of all polynomials with real coefficients, i.e.

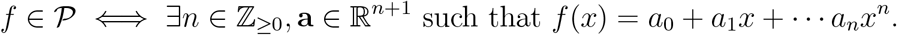

#### Definition 1.

Let 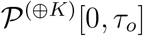 be the collection of piecewise polynomials with *K* internal breakpoints defined over [0, *τ_o_*]:

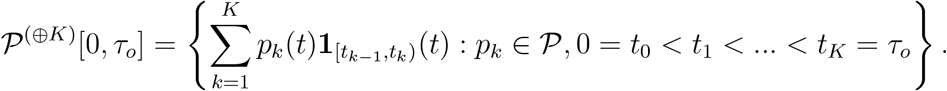

(The breakpoints *t*_0_,…, *t_K_* may vary between members of 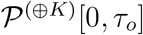.) Let 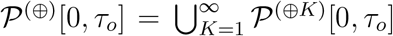 be the set of piecewise polynomials with any finite number of pieces. Similarly, we define the positive subset of these polynomials, 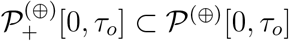, by

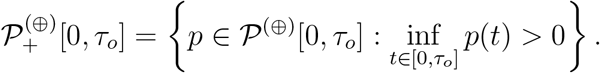

We now define the corresponding collection of phylogenetic BD models consisting of piecewise-polynomial birth and death rates. In the results that follow, we assume that *ρ* is known; if *ρ* is left to be estimated, then the resulting model space is unidentifiable even when λ and *μ* are constrained to be constant (Stadler, 2009; Stadler and Steel, 2019). Thus, the model class of interest supposes the sampling probability *ρ* is known in advance.

There are several equivalent ways to define the phylogenetic BD model. In this paper, we find it convenient to work in the following parameterization.

#### Definition 2.

Let

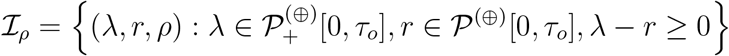

be the space of all piecewise-polynomial BD parameterizations with birth rates 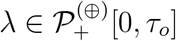, net diversification rates 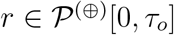, and fixed sampling fraction *ρ* ∈ (0, 1].

Note that elements of the model class 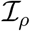 are in 1−1 correspondence with models defined by the more common birth/death rate parameterization, via the identity *r* = λ – *μ*. The function *r* is commonly referred to as the *net diversification rate* (Rabosky, 2010).

Louca and Pennell (2020) consider a new quantity called the *pulled speciation rate* λ_p_, defined as

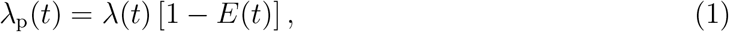

where *E* is the probability that a lineage that exists at time *t* is not sampled at the present. Thus, extinction “pulls” the observable birth rate downwards compared to the true one. *E* satisfies the differential equation

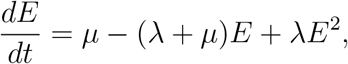

with initial condition *E*(0) = 1 – *ρ* (Morlon et al., 2011). This Bernoulli-type equation has the solution

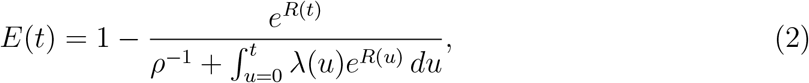

where

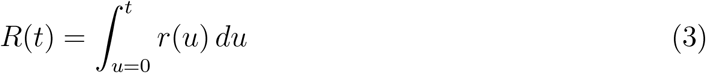

is the cumulative net speciation rate. Thus,

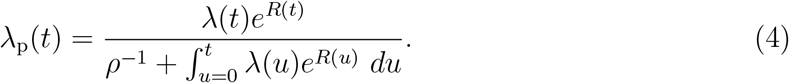

The cumulative integral Λ_p_ is given by

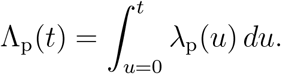

Finally, we write the likelihood of a timetree. For the phylogenetic BD process, the likelihood depends only on the number and timing of branching events, and is independent of the tree topology (Nee et al., 1994; Morlon et al., 2011). Given that the number of tips is *N* and the process survives to the present over a period of length *τ_o_*, (Louca and Pennell, 2020, supp. eqn. 34) show that the likelihood of a tree with bifurcation times *τ*_1_, …, *τ*_*N*–1_ is

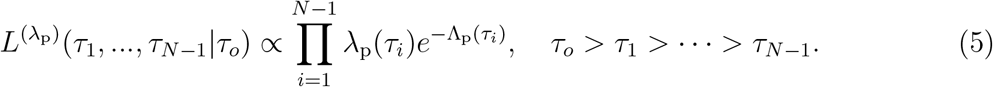

From the preceding display, it is clear that a sample of merger times (*τ*_1_, …, *τ*_*N*–1_) from an extant timetree can be equivalently viewed as the order statistics of *N* – 1 i.i.d. draws from a distribution with density

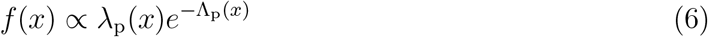

supported on [0, *τ_o_*], and that λ_p_ is the hazard rate function of this distribution. (See Section 5.2.1 for additional discussion.) Thus, λ_p_ completely characterizes the distribution of merger times in a timetree, and different BD models have different likelihoods if and only if their respective pulled rate functions are not equal almost everywhere on [0, *τ_o_*]. Moreover, subject to standard regularity conditions on hazard rate functions, λ_p_ (at least) can be consistently estimated from a timetree as the number of leaves *N* tends to infinity.

## 3 Results

We first show that piecewise polynomial phylogenetic birth-death models are identifiable from time-tree data, meaning that that different models in 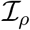 have different likelihood functions. As noted in the preceding section, it suffices to prove that different models possess different pulled rate functions.

#### Theorem 1.

(Identifiability of piecewise polynomial BD models). Let 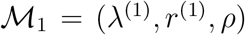 and 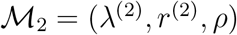 be two models in 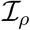. Then 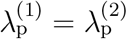 if and only if 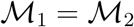.^1^

Theorem 1 is the main result of this paper, and should be contrasted with that of Louca and Pennell (2020). There, it is shown that, within the model class consisting of continuously differentiable birth and death rate functions, there are *infinitely* (in fact, uncountably) many pairs of such functions which all map to the same pulled rate function. Hence, statistical estimates of λ_p_, even if they were somehow uncontaminated by error, do not automatically translate into accurate estimates of the underlying rate functions. The best one can hope for is to estimate an equivalence or “congruence” class of rate functions, whose members can be qualitatively quite different, as Louca and Pennell exhibit. Theorem 1 asserts that this unfortunate situation cannot occur if one places additional and somewhat mild assumptions on the model class: different models in 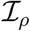 have different pulled rate functions, and conversely. In fact, the theorem shows that *every* unique piecewise-polynomial rate parameterization corresponds to a different pulled rate function. Hence, the number and placement of break points, as well as the underlying rates themselves, can, at least in principle, all be estimated given a sufficiently large time tree.

However, this seemingly positive outlook does not paint the full picture. In Section 4, we demonstrate that phylogenetic BD models can fail to be “practically” identifiable, in the sense that it may be impossible to ascertain the correct model given a realistic amount of data. To formalize this, we consider a testing problem where the task is to choose between two competing models which are hypothesized to have generated the data. One hypothesis is *H*_1_ : (λ, *r, ρ*), where now the rate functions λ and *r* can be arbitrary, and are not restricted in form as in Theorem 1. The second hypothesis differs from the first by only a multiplicative perturbation of the birth-rate function, *H*_2_ : ((1 + *η*)λ,*r, ρ*), where *η* > 0 is a constant. Intuitively, if *η* is too small relative to the amount of data that has been collected, then it is difficult to distinguish between *H*_1_ and *H*_2_. Our next result quantifies this intuition.

#### Theorem 2.

Consider the hypothesis testing problem where *H*_1_ states that the birth-death model over [0, ∞) is (λ, *r, ρ*) while *H*_2_ states that the birth-death model is ((1 + *η*)λ,*r, ρ*), where *η* > 0 is a constant. If an extant timetree on N leaves is observed, then for sufficiently small *η*, the Bayes error rate for distinguishing between *H*_1_ and *H*_2_ is at least (1 – ϒ)/2, where

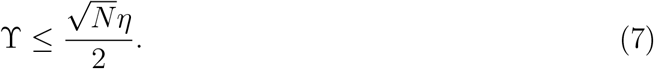

We prove a complementary result for when the net diversification rate *r* is scaled. However, for technical reasons, we are forced to make the strong assumption that λ and *r* are constant when proving this theorem.

#### Theorem 3.

Consider the hypothesis testing problem where *H*_1_ states that the birth-death model over [0, ∞) is (λ,*r, ρ*) while *H*_2_ states that the birth-death model is (λ, (1 + *η*)*r, ρ*), where both λ and *r* are constant over time, and *η* > 0 is a constant. If an extant time tree on *N* leaves is observed, the for sufficiently small *η*, the Bayes error rate for any classifier is at least (1 – ϒ)/2, where

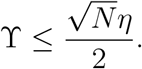

Since the Bayes error rate lower bounds classification accuracy, the theorems imply that the probability of correctly determining whether the timetree was generated by *H*_1_ or *H*_2_ is at most (1 + ϒ)/2 using *any* method. The key feature of the bounds is that they degrade in only the *square root* of the size of the time-tree *N*. Hence, unless *N* ≫ 1/*η*^2^, no procedure can distinguish between *H*_1_ and *H*_2_ with high probability.

## 4 Discussion

In this paper, we have derived new theoretical results concerning estimation of phylogenetic birth-death models using time-tree data. Our main result, Theorem 1, establishes identifiability of phylogenetic BD models parameterized by piecewise polynomial rate functions. One implication that may be surprising is that there exist identifiable classes of birth-death models with smoothly varying birth and death rates that are not purely constant. In particular, spline rate functions are identifiable. Splines are piecewise polynomials with additional smoothness constraints, and are widely used to model natural systems.

Theorem 1 improves on an earlier result of Legried and Terhorst (2022a), who established identifiability of piecewise constant BD models; we obtain their result as a special case. However, the sampling models underlying the two results are slightly different. Legried and Terhorst (2022a) assume access to a finite collection of *moments* of merger times (*τ*_1_,…, *τ_N_*), the number of which increases with the number of pieces *K* of the rate functions. Conceivably, these could be estimated using a large collection of independent time trees all of size *N*. Here, we assume access to the complete distribution of (*τ*_1_,…, *τ_N_*), which, as noted above, is given essentially by the density *f*(*x*) in equation (6). This scheme is more in keeping with the model studied by Louca and Pennell (2020), where there is a single time-tree tending in size to infinity.

To better understand the relationship between Theorem 1 and the non-identifiability result of Louca and Pennell, consider Figure 1. The top row is adapted from Figure 1 of their paper, and depicts four pairs of (λ, *μ*) rate functions which all have the pulled sampling rate λ_p_. In the bottom row, we approximated each of these functions using a cubic spline, with sixteen knots placed at *t* = 15, …, 0. The models shown in the bottom row are identifiable; the ones in the top are not. Note that there are some visual differences between the two panels; for example the speciation rates of the rate and green models intersect in the top row, whereas the spline smoothness constraints prevent them from doing so in the bottom. (For illustrative purposes, we used a very simple interpolating spline with equispaced knots; a closer approximation could be obtained using a more complex fitting procedure.) Practitioners must decide if the spline model class (or a related class of identifiable models) can faithfully model population dynamics in their application.

**Figure 1:**
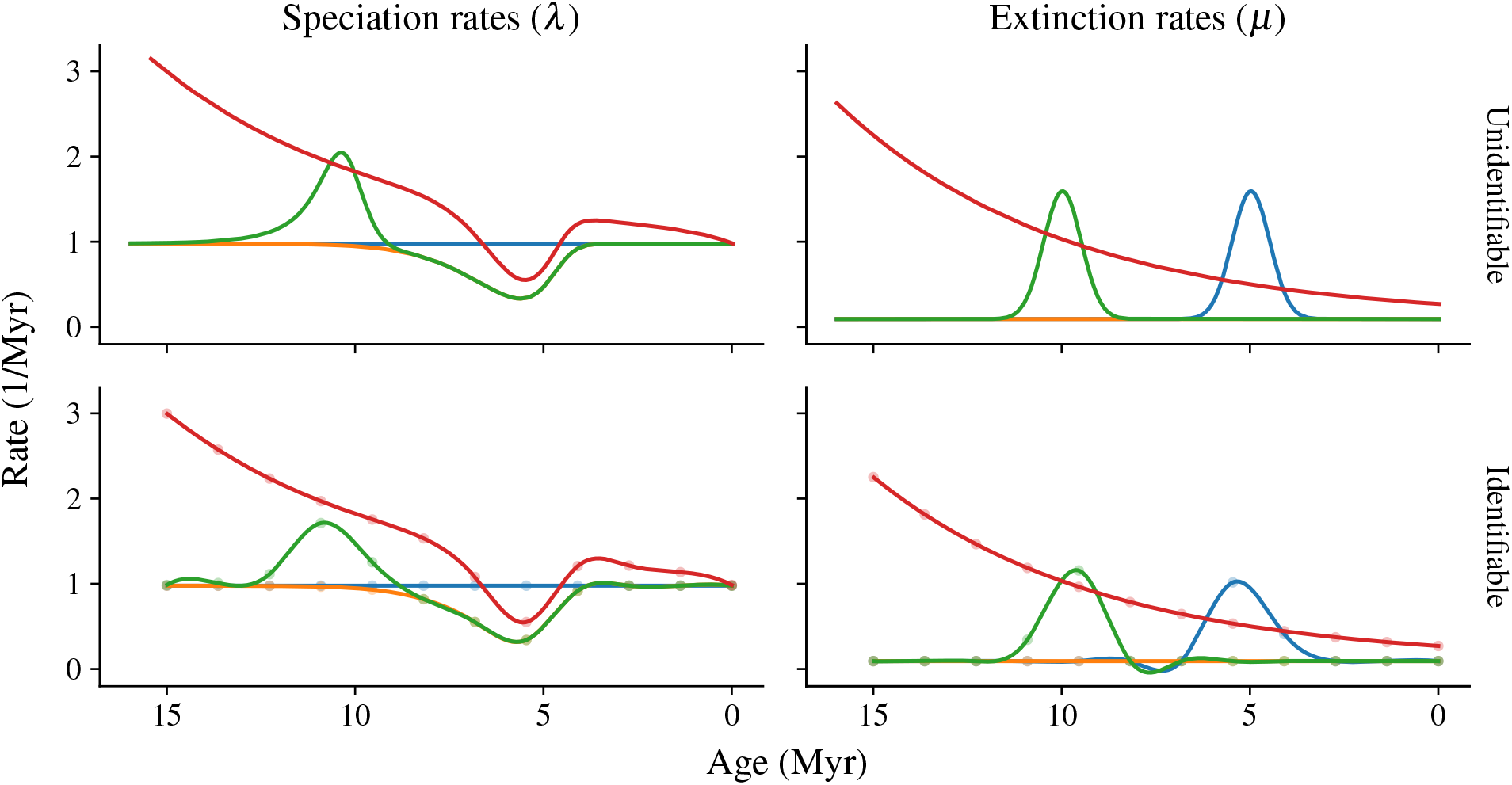
Identifiable versus unidentifiable models. The top row contains four unidentifiable models shown in Figure 1 of Louca and Pennell (2020). Each color-coded (λ, *μ*) pair in the top maps to the same pulled rate function; hence they are indistinguishable. In the bottom row, we approximated these functions using interpolating cubic splines, with knots at *t* = 15, 14,…, 0. According to Theorem 1, these models have different pulled rate functions, and may be distinguished in the infinite-data limit.

The results we present here restore to some extent the mathematical footing beneath the many published studies that have utilized the linear birth-death process to describe evolution. However, identifiability is a minimal regularity condition one can impose on a statistical model, and estimation remains challenging. To see this, consider now Figure 2, which plots the pulled speciation and net diversification rates for each of the spline-based models from Figure 1. By Theorem 1, these functions are necessarily different—but in terms of estimation, the important question is *how* different they are. It is evident from Figure 2 that distinguishing the green model from the other three is probably feasible. In contrast, the red and orange models appear almost identical in terms of λ_p_, and identifying which of them generated a particular data set is likely to be difficult.

**Figure 2:**
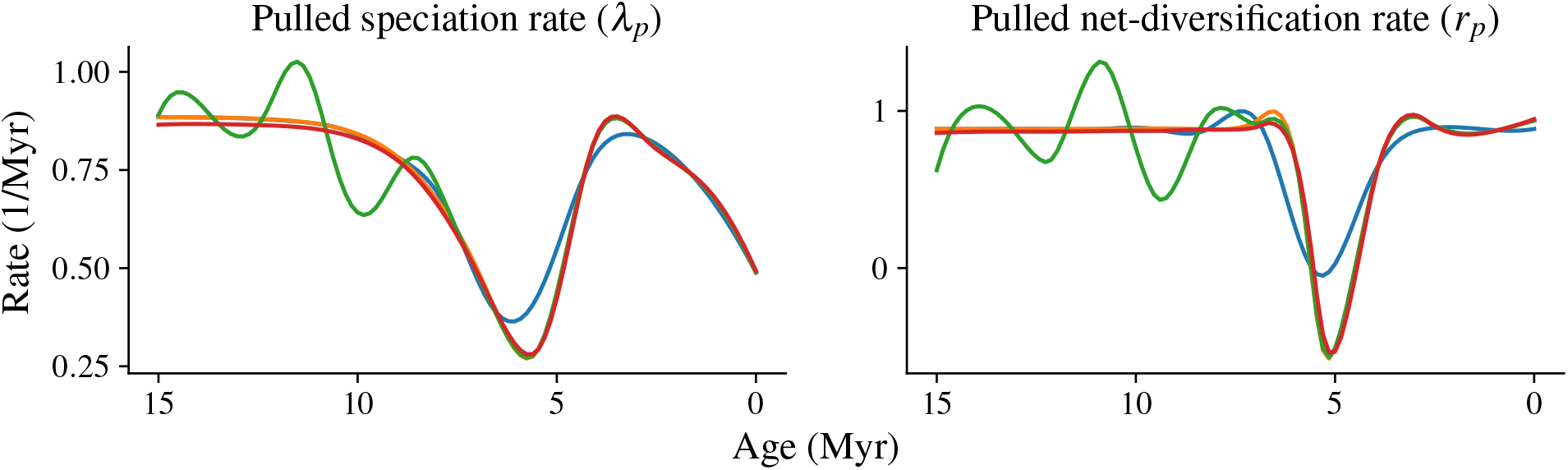
The pulled rates of speciation and net diversification for each of the spline models shown in the bottom row of Figure 1. The sampling fraction was *ρ* = 0.5 in each model.

This leads to our second set of results, Theorems 2 and 3, which study the hardness of distinguishing between competing BD hypothesis using only a finite amount of data. We prove that even answering the relatively simple question of whether the data are generated by a particular model, or one in which its rate function(s) are a scalar multiple of it, scales poorly in the time-tree size *N*. (This theoretical limitation is also found in coalescent models, see Kim et al., 2015.) As Figure 2 already suggests, the question of when practical estimation is possible is likely to be quite subtle even in identifiable model classes. Much more deserves to be said, and this is an important area for future research.

Finally, an obvious caveat to the results we have presented is that it may not be possible to estimate λ_p_ in the first place. As noted in Section 1, inferring λ_p_ is tantamount to estimating the hazard rate function of the distribution shown in (5). If it were possible to directly sample from this distribution, estimation of λ_p_ would be routine, however this is not possible in practice. Rather, one must first estimate the time-tree itself, and then treat the estimated merger times as though they were samples from (5). Tree inference is itself a difficult problem, and often these estimates contain considerable error, so it is not obvious that such a procedure leads to accurate downstream estimates of the underlying BD model, even in the infinite-data limit.

The assumption that λ_p_ is known colors our results as follows. First, it implies that the lower bounds derived in Theorems 2 and 3 are likely not sharp, even for identifiable model classes, and that inferring phylogenetic BD models can be even harder than is indicated by theorems. Conversely, it qualifies Theorem 1 somewhat, since the theorem does not resolve the question of whether phylogenetic BD models are identifiable *on the basis of the observed data*. In this sense, unidentifiability results like those derived in Stadler (2009); Louca and Pennell (2020) are stronger, since they imply that the model cannot be estimated *even if* we somehow had direct access to the underlying time-tree. We note in closing that there is a growing literature on phylogenetic identifiability (e.g., Rhodes and Sullivant,2012; Mossel and Roch, 2013), which derives conditions under which it is possible to consistently estimate an underlying phylogeny given character data evolving along its branches. If it can be shown that an asymptotically expanding time tree of the variety considered here is estimable, it would, together with our results, imply identifiability of polynomial rate functions from character data.

## 5 Proofs

This section contains the proofs of Theorems 1, 2 and 3.

### 5.1 Polynomial identifiability

In this subsection, we prove Theorem 1. We begin with the case of polynomial models.

#### Lemma 1.

Consider two models (λ^(1)^, *r*^(1)^, *㋁*) and (λ^(2)^, *r*^(2)^, *ρ*) where 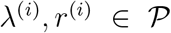 are polynomials with inf_*t*∈[0,*τ_o_*_] λ^(*i*)^ (*t*) > 0, and define

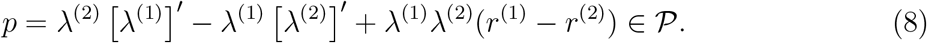

If *u* is a limit point of the set 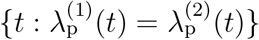, then *p*(*u*) = 0. In particular, if there are infinitely many such points, then *p* ≡ 0.

*Proof*. For any such *u*, we have by continuity

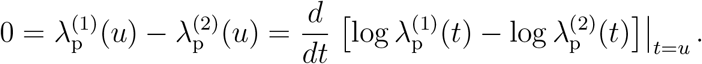

By (4),

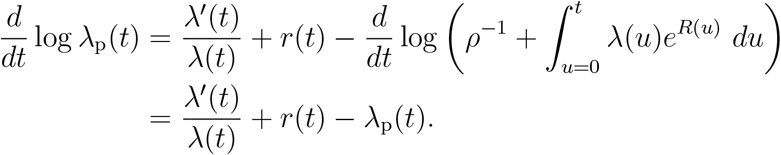

Thus

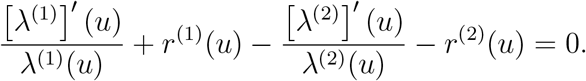

Clearing denominators gives *p*(*u*) = 0. If there are infinitely many limit points, then *p* has infinitely many zeros and hence is identically zero.

Our main new technical result is the following sufficient condition for equality of two polynomial models, which works by an algebraic argument.

#### Lemma 2.

Let (λ^(1)^, *r*^(1)^, *ρ*), (λ^(2)^, *r*^(2)^, *ρ*), and *p* be as defined in Lemma 1. If 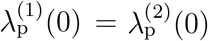 and *p* ≡ 0, then (λ^(1)^, *r*^(1)^) = (λ^(2)^, *r*^(2)^).

*Proof*. By assumption, for *i* ∈ {1,2} there exist *m_i_* = deg λ^(*i*)^ and *n_i_* = deg *r*^(*i*)^ and real coefficients 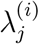 and 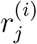, with 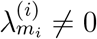 such that

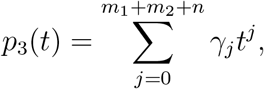

We first establish *r*^(1)^ = *r*^(2)^. Define *n* = max(*n*_1_, *n*_2_). If *n* = –∞ then *r*^(1)^ = *r*^(2)^ ≡ 0 and there is nothing to show. Otherwise, let

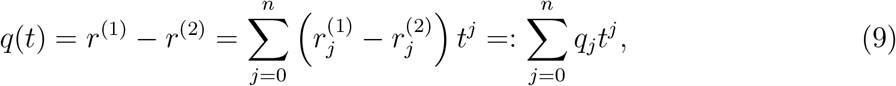

where we set 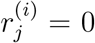 for all *j* > *n_i_*. In order to prove *r*^(1)^ = *r*^(2)^, it suffices to show *q_j_* = 0 for all *j* ∈ {0,…, *n*}.

Next, define the polynomials

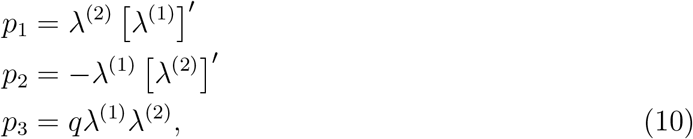

so that

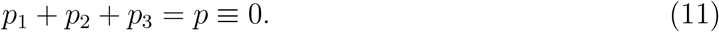

We observe that deg(*p*_1_ + *p*_2_) ≤ *m*_1_ + *m*_2_ – 1, and that *p*_3_ is a polynomial of degree at most *m*_1_ + *m*_2_ + *n*:

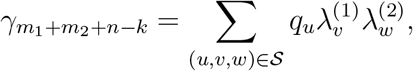

for some 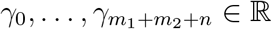. Hence, by (11),

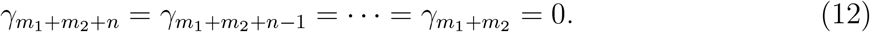

We proceed to establish that *q*_*n*–*j*_ = 0 for 0 ≤ *j* ≤ *n* by induction on *j*. For the base case *j* = 0, we have by (10) that the leading coefficient of *p*_3_ equals the product of the leading coefficients of λ^(1)^, λ^(2)^, and *q*:

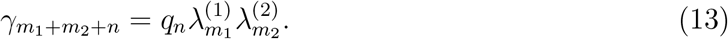

Since 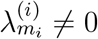, it must be that *q_n_* = 0.

Now suppose the claim is true for all 0 ≤ *j* < *k*. By polynomial convolution, we have

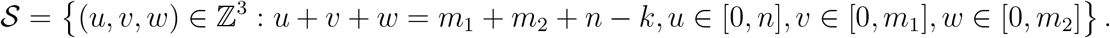

where the summation is over the index set

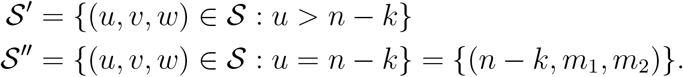

We partition 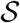 into disjoint subsets

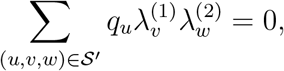

By the inductive hypothesis, so that

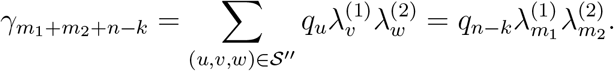

This together with (12) implies that *q*_*n*–*k*_ = 0. Hence *q* = 0, so *r*^(1)^ = *r*^(2)^.

We conclude by showing that λ^(1)^ = λ^(2)^. Since *q* = 0, we have *p*_1_ = –*p*_2_ by (11).

Therefore,

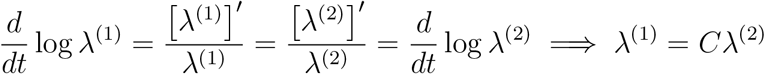

for some constant *C* > 0. Hence,

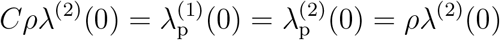

implies *C* = 1.

The lemma furnishes the following corollary on identifiability for polynomial models.

#### Proposition 1.

Let (λ^(1)^, *r*^(1)^, *ρ*) and (λ^(2)^, *r*^(2)^, *ρ*) be as defined in Lemma 1. If (λ^(1)^, *r*^(1)^) ≠ (λ^(2)^, *r*^(2)^), then 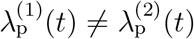 on a subset of positive measure in [0, *τ_o_*].

*Proof*. By Lemma 2, either *p* ≠ 0 or 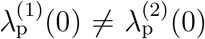. In the former case, Lemma 1 shows that set 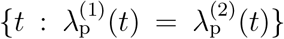 contains at most a finite number of limit points, so it is null. In the latter case, by continuity, there exists *u* > 0 such that 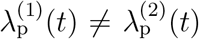 for all *t* ∈ [0, *u*).

We use Proposition 1 to prove Theorem 1 for piecewise-polynomial birth and death rates on an arbitrary number of pieces. The proof is an extension of Proposition A.5 in Legried and Terhorst (2022a).

#### Proof of Theorem 1.

The “if” direction is immediate. To establish the “only if” direction, we show its contrapositive: (λ^(1)^, *r*^(1)^) = (λ^(2)^, *r*^(2)^) implies 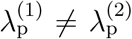 on a set of positive measure. We may assume that the two models are defined on the same set of breakpoints, 0 = *t*_0_ < … < *t_K_* = *τ_o_*, since this can always be achieved by increasing *K*. Then there exists a non-empty interval [*u,v*) ⊂ [0, *τ_o_*] such that

1. (λ^(1)^(*s*), *r*^(1)^(*s*)) = (λ^(2)^(*s*), *r*^(2)^(*s*)) for all 0 < *s* < *u*;
2. (λ^(1)^(*s*), *r*^(1)^(*s*)) = (λ^(2)^(*s*), *r*^(2)^(*s*)) for all *s* ∈ [*u,v*); and
3. [*u,v*) ⊂ [*t_k_, t*_*k*+1_) for some *k*.

(We could have *u* = 0, rendering the first condition vacuous.) Let the birth and net diversification rates for the two models over [*u,v*) be denoted 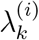 and 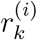, respectively.

Recall the function *E*(*t*) defined in equation (2). Because *E* is continuous and *E*^(1)^(0) = *E*^(2)^(0) = 1 – *ρ*, condition (1) above implies that *E*^(1)^(*u*) = *E*^(2)^(*u*) = 1 – *ε* ∈ (0, 1) for some *ε*. Define the shifted models (*π*^(*i*)^, *q*^(*i*)^, ∈) for *i* ∈{1, 2}, where

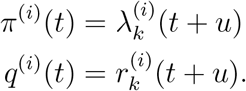

Denote the corresponding pulled rate functions 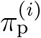, so that 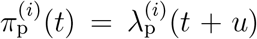 for *t* ∈ [0, *v* – *u*). By Proposition 1, 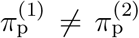 on a non-null subset of [0, *v* – *u*). The same is therefore true of 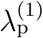 and 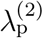 over [*u,v*).

### 5.2 Lower bounds

In this section, we prove Theorems 2 and 3. Subsections 5.2.1 and 5.2.2 contain some necessary background and definitions, and can be omitted if the reader is already familiar with them. The results of this section use techniques developed in Kim et al. (2015); Legried and Terhorst (2022b).

#### 5.2.1 The pulled speciation rate as a hazard rate

Let the origin time *τ_o_* have an (improper) uniform prior distribution on (0, ∞). Then the likelihood of *τ_o_* > *τ*_1_ > … > *τ*_*N*–1_ is given by

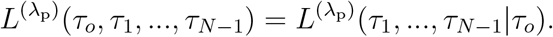

The likelihood formula holds, in particular, when we take *τ_o_* → ∞. Then for any sequence *τ*_1_ > *τ*_2_ > … > *τ*_*N*–1_ > *τ_N_* = 0, the likelihood is

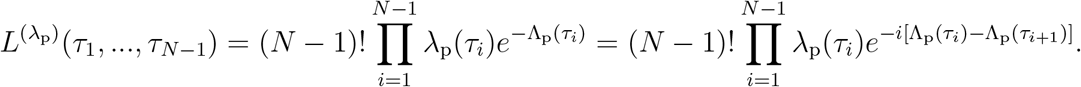

This shows that λ_p_ is a hazard rate function. To ensure that the integral of the likelihood function converges, we assume that

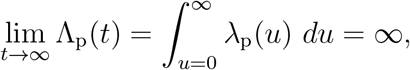

so this requirement is satisfied. Since λ_p_ is a hazard rate, it corresponds to a probability density function *f_p_*(*t*) = λ_p_(*t*)*e*^−Λ_p_(*t*)^.

#### 5.2.2 Bounding the Hellinger distance

Here, we recall the total variation distance and Hellinger distance between measures, borrowing from the explanation in Legried and Terhorst (2022b). Consider a measurable space 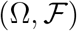 with two possible probability measures *P* and *Q* on it. The two probability spaces 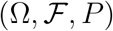 and 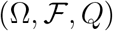 then have corresponding probability density functions *f_P_* and *f_Q_*. The *total variation distance* between *P* and *Q* is defined as

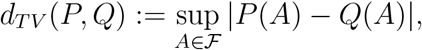

which equals 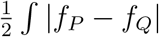. As this integral is itself a metric, one might abuse notation and write *d_TV_*(*f_P_, f_Q_*) to mean the same thing.

Suppose a datum *D* has been generated under *P* or *Q* and we want to decide which measure was used, assuming both choices are equally likely. The total variation distance between *P* and *Q* bounds the ability of any classifier to decide correctly. Let *χ* ∈ {*P, Q*} be the true data generating distribution, and let 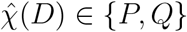 be a classifier. The probability that 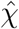 correctly classifies *D*, i.e. 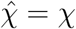, is

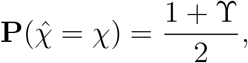

where ϒ > 0 because the error of any binary classifier can be made less than 1/2. Another way to write ϒ is

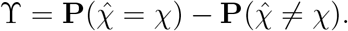

It can be shown that the best possible classification rule is the likelihood ratio: 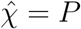 if and only if *P*(*D*) > *Q*(*D*). In that case, we have

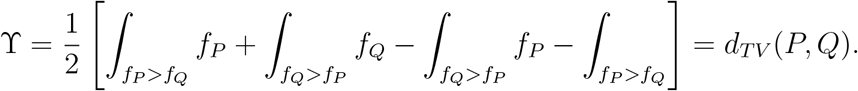

The likelihood ratio is said to achieve the *minimal error rate* or *Bayes error rate*. If multiple independent samples *D*_1_,…, *D_L_* are available, then similarly ϒ equals *d_TV_*(*P*^⊗*L*^, *Q*^⊗*L*^), where *P*^⊗*L*^ denotes product measure.

In our problem setting, we find it easier to work with the Hellinger distance, which is given defined as

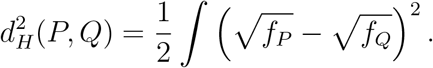

The Hellinger distance is related to total variation via

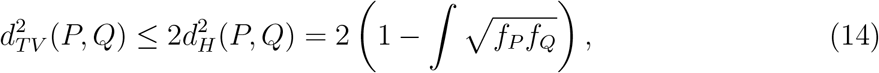

and it also possess the following subadditivity property: if 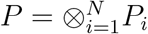 and 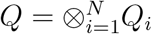 are product measures, then

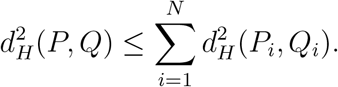

By bounding the right-hand side above by a quantity converging to 0 quickly, it follows that the total variation distance also converges to 0 quickly.

We will prove Theorems 2 and 3 by computing the Hellinger distance of competing joint distributions of (*τ*_1_, …, *τ*_*N*–1_). For this, we need to establish precise two-sided bounds of 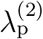 in terms of 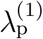. We start in the setting of Theorem 2, where λ is the birth rate corresponding to
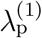 and (1 + *η*)λ is the birth rate corresponding to 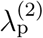. The two-sided bound of 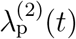 is expressed in the following Lemma.

#### Lemma 3.

Let *t* > 0 and *η* > 0 be arbitrary. Then

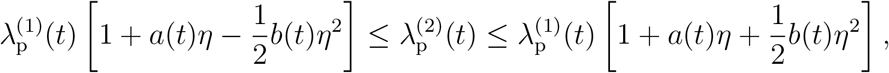

where

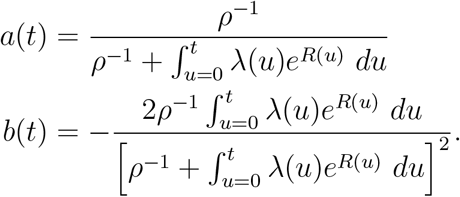

*Proof*. The quotient is

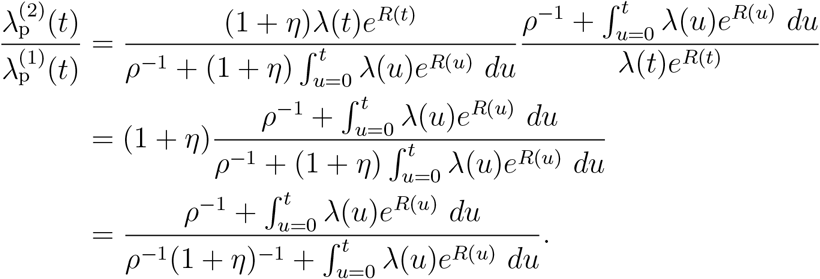

The coefficients *a*(*t*) and *b*(*t*) are obtained by differentiating the above expression with respect to *η* and evaluating at 0. The first derivative is

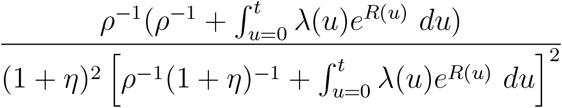

which evaluates at 0 to

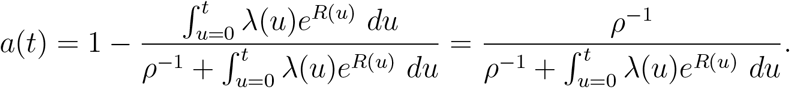

Note that the right-hand side is a positive number between 0 and 1. The second derivative is

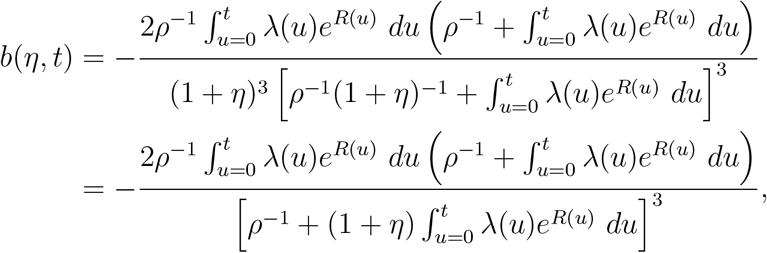

which is always negative. We now apply Taylor’s theorem, observing that

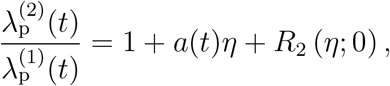

where the remainder term *R*_2_(*η*; 0) is

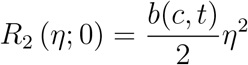

for some *c* ∈ (0, *η*). Since *∂*|*b*(*c, t*)|/*∂c* < 0, we have |*R*_2_(*η*;0)| < *b*(*t*)*η*^2^/2, where

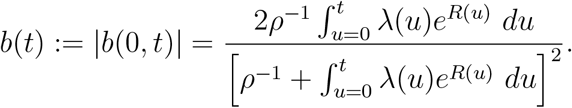

Then

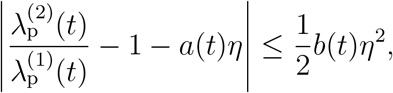

which implies the result.

To use the Lemma to prove Theorem 2, we show that *a*(*t*) and *b*(*t*) are each bounded by constants over *t* ≥ 0.

#### Proof of Theorem 2.

From (14), the squared error rate satisfies

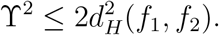

Starting with the case of *N* = 2, we have 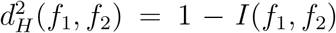 where 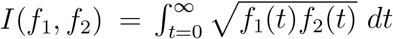. Defining the probability density function

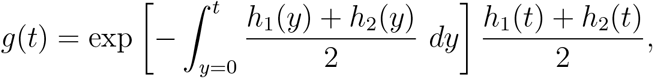

we have

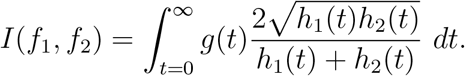

Now we use an upper bound on *h*_1_ + *h*_2_ and a lower bound on *h*_1_*h*_2_ from Lemma 3 to obtain a lower bound for 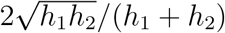. We have

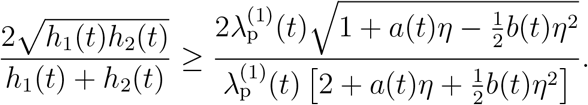

The power series expansion of the right-hand side is

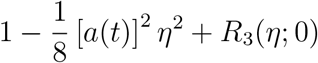

with the remainder term satisfying

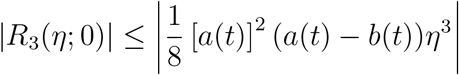

for some *t* ∈ [0, *η*). Both functions *a*(*t*) and *b*(*t*) are bounded in *t*. So for *η* close enough to 0, independently of *t*, we have

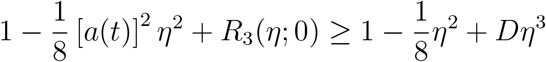

where *D* is a non-zero constant independent of *t*. So since *g*(*t*) is a density,

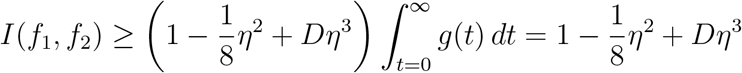

for *η* sufficiently close to 0. Putting it together, for such *η* the Hellinger distance is bounded as

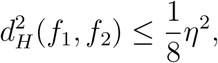

and the Theorem follows in the *N* = 2 case.

Now we derive the bound for an arbitrary number *N* of extant individuals. There are *N* – 1 independent coalescent times with identically distributed arrival times. As *f*_1_ and *f*_2_ are product measures of *N* – 1 independent random variables, the subadditivity property implies

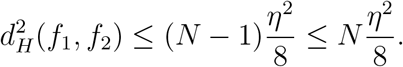

Then Theorem 2 follows from (14).

For Theorem 3, we compare hypothetical models with the same birth rate and sampling probability, but *R*^(1)^(*t*) = *R*(*t*) and *R*^(2)^(*t*) = (1 + *η*)*R*^(1)^(*t*) for all *t*. Although we prove Theorem 3 in the case where *R*^(1)^ is constant over time, the analogue to Lemma 3 can be proved for general λ^(1)^ and *R*^(1)^.

#### Lemma 4.

Let *t* be fixed and *η* be a positive number close to 0. Then

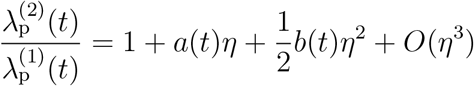

as *η* → 0, where

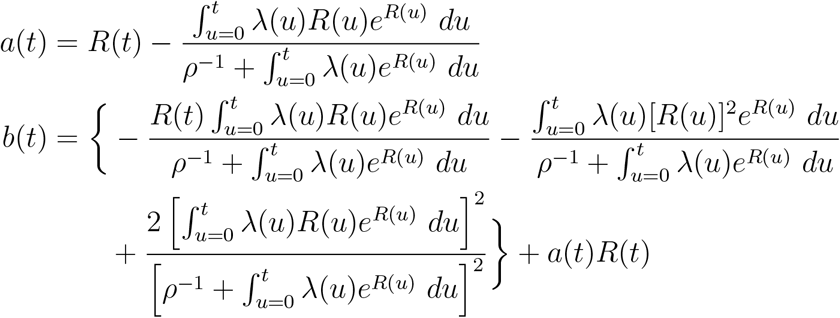

*Proof*. The quotient is

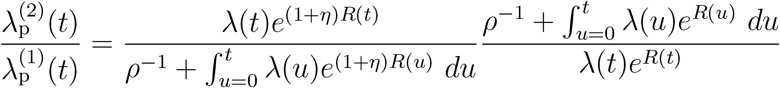

As before, the coefficients *a*(*t*) and *b*(*t*) are obtained by differentiating the above expression with respect to *η* and evaluating at 0. The first derivative is

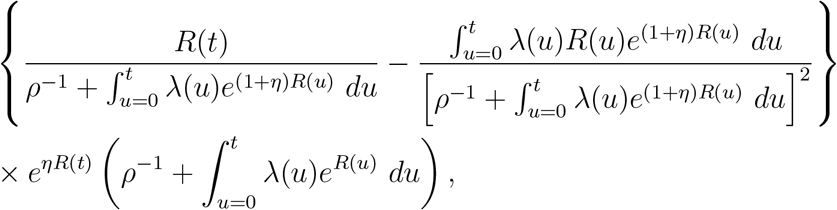

which evaluates at *η* = 0 to

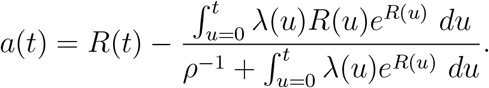

The second derivative is

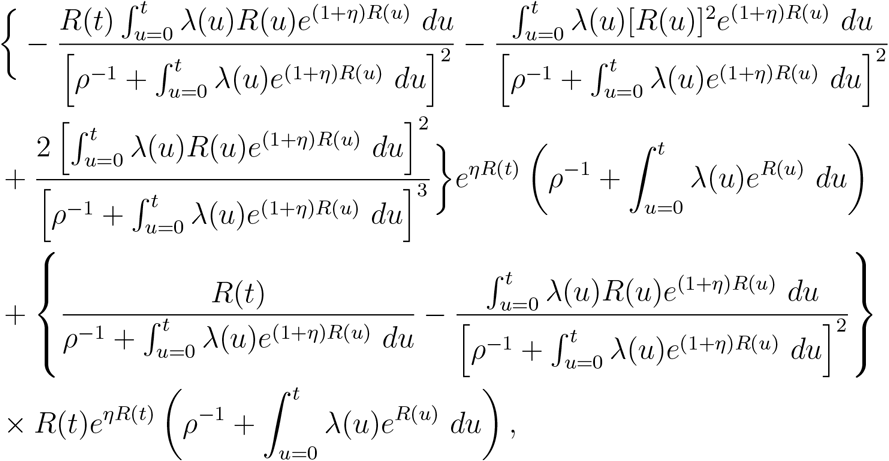

which evaluates at *η* = 0 to

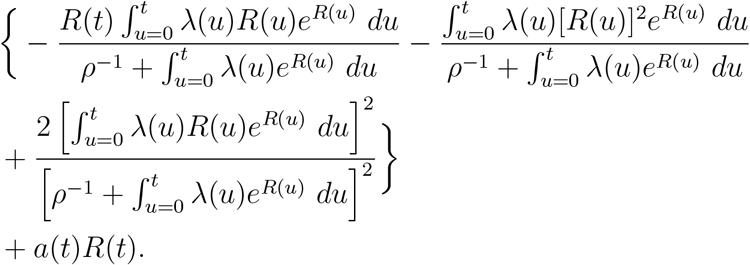

The proof of Theorem 3 is similar. However, because the *R*(*t*) function appears inside exponents, bounding *a*(*t*) and *b*(*t*) is more complicated. This is why we assume that *R*(*t*) = *Rt* is linear in the theorem.

#### Proof of Theorem 3.

Going forward, we assume that the rates are constant, implying there exists a real number *R* such that *R*(*t*) = *Rt* for all *t*. Then we use Taylor’s theorem:

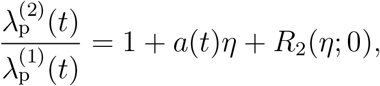

where now

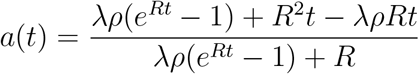

by Lemma 4. If *R* > 0, then |*a*(*t*)| ≤ 1 for all *t* ≥ 0. If *R* < 0, then the limit of *a*(*t*) as *t* → +∞ is –∞, which is not usable. So we proceed with *R* > 0. The remainder term *R*_2_(*η*;0) has the same structure as before, but

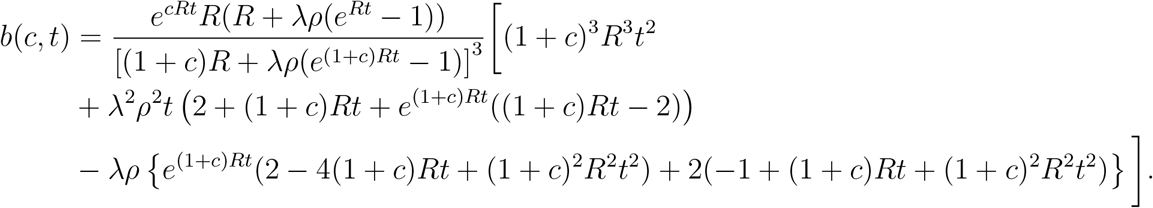

(As *R* > 0, there is no possibility of division by 0.) Now we derive c-independent bounds for the bracketed expression. The negative terms are dropped and the remaining polynomial terms are replaced with exponentials:

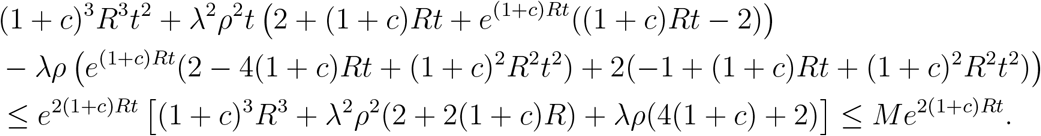

On the right-hand side, the factor *M* can be taken to depend only on λ as *ρ* ≤ 1, *R* ≤ λ, and *c* ∈ (0, *η*] ⊂ (0, 1). The companion lower bound for this polynomial

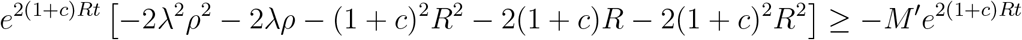

is for a factor *M*′ depending only on λ. Going back to *b*(*c, t*), we drop the negative term in the numerator to get

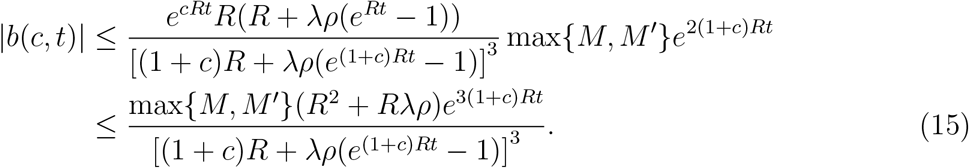

The proof is completed by using Lemma 5, below, to bound |*b*(*c, t*)|. By the lemma, we have 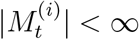 for the functions 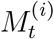 defined in the statement below. Therefore, |*b*(*c, t*)| is bounded above by some constant independent of *c* and *t*. So *b*(*t*) = *b*(0, *t*) is bounded in t. Since *a*(*t*) and *b*(*t*) are bounded in *t*, we can prove the analogous result for scaling *R*. This is sufficient to ensure 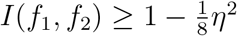 as in Theorem 2.

#### Lemma 5.

Letting *M*″ = max{*M, M*′}, we have 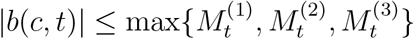 where

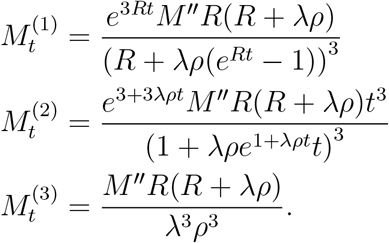

*Proof*. In the two-sided bound (15), we optimize over *c* ∈ (0, ∞) for a fixed *t*. The partial derivative in *c* is

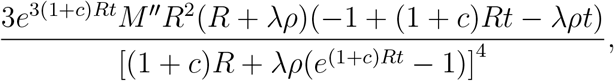

yielding the critical point

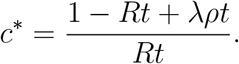

Since the function is bounded, the global maximum is obtained either at a critical point or an end point of the interval *c* ∈ [0, ∞). Evaluating the original bound at *c* = 0 and *c*\* and taking the limit *c* →+∞ yields 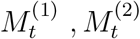, and 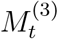, respectively.

## Acknowledgments

This research was supported by NSF grants DMS-2052653 and DMS-1646108.

## Footnotes

1 Throughout the paper, we take *f* = *g* to mean that the rate functions *f* and *g* are equal almost everywhere with respect to Lebesgue measure.

